# Mechanical coupling with the nuclear envelope shapes the *S. pombe* mitotic spindle

**DOI:** 10.1101/2022.12.28.522145

**Authors:** Marcus A Begley, Christian Pagán Medina, Taylor Couture, Parsa Zareiesfandabadi, Matthew B Rapp, Mastawal Tirfe, Sharonda LeBlanc, Meredith D. Betterton, Mary Williard Elting

**Affiliations:** Department of Physics, North Carolina State University, Raleigh, NC; Department of Biology, Duke University, Durham, NC; Department of Physics and Molecular, Cellular, and Developmental Biology, University of Colorado, Boulder, CO; Quantitative and Computational Developmental Biology Cluster, North Carolina State University, Raleigh, NC

## Abstract

The fission yeast *S. pombe* divides via closed mitosis, meaning that spindle elongation and chromosome segregation transpire entirely within the complete nuclear envelope. Both the spindle and nuclear envelope must undergo significant conformation changes and exert varying forces on each other during this process. Previous work has demonstrated that nuclear envelope expansion^1,2^ and spindle pole body (SPB) embedding in the nuclear envelope^3^ are required for normal *S. pombe* mitosis, and mechanical modeling has described potential contributions of the spindle to nuclear morphology^4,5^. However, it is not yet fully clear how and to what extent the nuclear envelope and mitotic spindle each directly shape each other during closed mitosis. Here, we investigate this relationship by observing the behaviors of spindles and nuclei in live mitotic fission yeast following laser ablation. First, we characterize these dynamics in molecularly typical *S. pombe* spindles, finding them to be stabilized by dense crosslinking, before demonstrating that the compressive force acting on the spindle poles is higher in mitotic cells with greater nuclear envelope tension and that spindle compression can be relieved by lessening nuclear envelope tension via laser ablation. Finally, we use a quantitative model to interpret how these data directly demonstrate that fission yeast spindles and nuclear envelopes are a mechanical pair that can each shape the other’s morphology.

## RESULTS AND DISCUSSION

### *S. pombe* mitotic spindles are highly crosslinked

The *Schizosaccharomyces pombe* mitotic spindle consists of a single bundle of 10-20 microtubules, held together along their lengths by microtubule-crosslinking proteins^6–8^. In the spindle midzone, microtubules of alternating geometric polarity form a square lattice^6^. During anaphase, motor proteins can crosslink and slide apart antiparallel microtubule neighbors, creating extensile force within the spindle and driving spindle elongation^7,9^. In *S. pombe*, this elongation is quite dramatic, with spindles elongating from hundreds of nanometers in length to roughly 10 μm over the course of mitosis^10,11^. Accompanying spindle elongation, the closed nuclear envelope undergoes a series of shape transitions, from a spheroid, through a peanut-shaped intermediate, to a dumbbell that will ultimately separate into two daughter nuclei (Figure 1A). Furthermore, both spindle elongation and nuclear shape are quite spatiotemporally stereotyped, adhering to a strict protocol throughout mitosis^12^, suggesting that fission yeasts can precisely tune their mitotic forces. The simplicity and standardization of the *S. pombe* mitotic apparatus allows us to effectively probe their fundamental physical characteristics by subjecting them to different molecular perturbations. We began by characterizing spindles in control *S. pombe* cells using laser ablation of live cells (Figure 1B)^13^. Upon ablation, the two spindle halves have previously been shown to collapse toward each other in a motor driven process^9,13,14^, bringing the two poles toward each other and reforming a single microtubule bundle (Figure 1B and Supplemental Video SV1). Before each spindle reforms, the two half-spindles are detached from each other, and microtubules within each half bundle can either stay tightly associated with each other, retaining their status as a single bundle, or become detached along their lengths, a process that we term “splaying”^15^.

**Figure 1.**
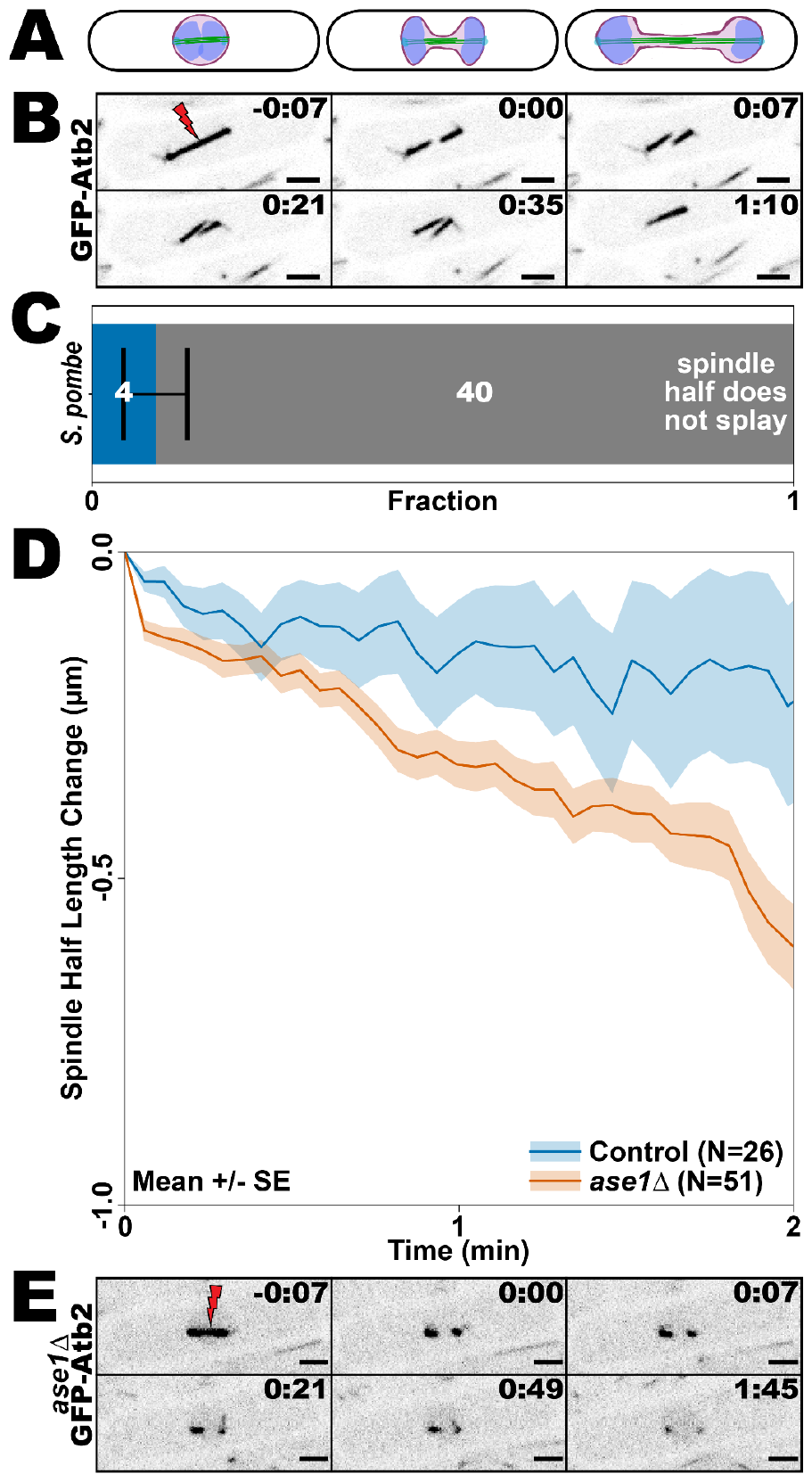
*S. pombe* mitotic spindles are highly crosslinked by Ase1, which stabilizes microtubules against depolymerization. All scale bars, 2 μm, timestamps, min:sec. (A) Cartoon showing anaphase nuclear deformation and spindle elongation in a *S. pombe* cell. (B) Typical example of laser ablation of an *S. pombe* spindle expressing GFP-Atb2 (MWE2), showing post-ablation spindle collapse, followed by the reattachment of ablated spindle halves at their plus-ends. (C) Ablated *S. pombe* spindle halves rarely splay apart. Error bar indicates the square root of the number of splayed events, as an estimate on the variation in this number assuming Poisson statistics. (D) *ase1Δ S. pombe* (orange) halves tend to depolymerize more quickly and severely than those from *S. pombe* with unperturbed ase1 (blue), as shown by traces of the average change in spindle half length after ablation. Traces represent average across all videos and shaded regions represent average +/-standard error on the mean. (E) Typical example of spindle ablation in *ase1Δ S. pombe* expressing GFP-Atb2 (MWE3), in which both spindle halves depolymerize. Some spindle splaying is also visible during depolymerization (0:49).

Comparison of the probability of each of these two behaviors is an indicator of the degree of crosslinking between the microtubules in the ablated spindle^15^. In *S. pombe*, ablated spindle halves splay only rarely (Figure 1C), suggesting a high degree of crosslinking, consistent with previous work characterizing *S. pombe* spindle structure^6,11^. Another striking feature of the behavior of *S. pombe* spindle halves following severing by laser ablation is that their overall length is maintained, with very little shortening (Figure 1D). The spindle at the point of severing (generally near the center of the spindle) forms an antiparallel architecture^6^, and thus we expect about half of the microtubule ends in each ablated half spindle to have their plus-ends facing toward the site of ablation. A typical hallmark of laser ablated microtubules in many cell types is depolymerization of exposed plus-ends^16–20^. Thus, it was initially surprising to us that such behavior is so rarely observed in these spindles (Figure 1B, D). High densities of crosslinking proteins have been shown to stabilize microtubule bundles against catastrophic depolymerization^21,22^. Therefore, the fact that spindle halves mostly resist depolymerization following ablation provides additional evidence that *S. pombe* spindles are very highly crosslinked. Indeed, such crosslinking is also consistent with the highly stereotyped organization of the central spindle at this stage^6^. Altogether, these data paint a picture of the *S. pombe* mitotic spindle as a tightly-bound microtubule bundle that is stable, even under significant physical disturbance.

### Ase1 crosslinking prevents microtubule depolymerization in *S. pombe*

An important component of the *S. pombe* mitotic spindle is the passive microtubule-microtubule crosslinking protein Ase1, which preferentially localizes to antiparallel microtubule overlaps in the spindle midzone, particularly at anaphase^11,23–25^.There, it functions to stabilize the spindle midzone, supports bipolar spindle formation in *S. pombe*, and even allows the formation of a bipolar spindle in the absence of kinesin-5 mediated antiparallel microtubule sliding^26^. Intriguingly, budding yeast Ase1 has also been shown, through the same underlying mechanism of stabilizing antiparallel overlaps, to slow motor-driven anaphase *in vivo*^*8,27*^ and sliding *in vitro*^*8,27*^. Thus, we sought to investigate the role of Ase1 in *S. pombe* spindle crosslinking.

To this end, we laser ablated mitotic spindles in ase1-deletion (*ase1Δ*) *S. pombe*. Strikingly, and in contrast to results above with normal Ase1 expression, rapid depolymerization of ablated spindle halves was common in *ase1Δ* cells (Figure 1D, E and Supplemental Video SV2), consistent with the role of Ase1 in stabilizing bundles^21,26,27^. Though this depolymerization prevented reliable characterization of post-ablation spindle half splaying as for control *S. pombe* cells (Figure 1C and Supplemental Videos SV1, SV2), we anecdotally observed some splaying of the remaining spindle halves as they depolymerized (Figure 1E), a behavior that was unusual in control cells (Figure 1C). These data are consistent overall with a central role for Ase1 in both crosslinking *S. pombe* spindles and stabilizing their microtubules against depolymerization.

### Increased nuclear envelope tension can bend spindles and slow their elongation

We next investigated the mechanical relationship between the *S. pombe* spindle and nuclear envelope. During spindle elongation, the spindle and nuclear envelope are mechanically linked at the two spindle pole bodies (SPBs), which nest into the nuclear envelope during mitosis in *S. pombe*^*28*^. Presumably, this connection allows nuclear envelope tension to be transmitted to the spindle^1^ and vice versa, but we sought to probe this mechanical relationship directly and quantitatively. To do so, we altered the nuclear envelope with cerulenin, a fatty acid synthesis inhibitor^29–31^ that prevents the addition of phospholipids to the nuclear envelope during nuclear envelope expansion, and has previously been shown to alter *S. pombe* mitotic spindle and nuclear shape^1,32,33^. We first quantified the behaviors of spindles in cerulenin-treated cells, and then performed laser ablations of both the spindle and nuclear envelope in these cells.

As has previously been observed in *S. pombe*^1^, we find that cerulenin treatment often causes bending in anaphase spindles, presumably due to increased nuclear envelope tension (Figure 2A, Supplemental Video SV3). Here, we quantify this bending, which we term ‘buckling,’ as a function of spindle length, and find that the onset occurs fairly abruptly when spindles reach a length of ∼4-5 μm (Figure 2B). In comparison, spindles in control cells rarely show significant curvature. (While Fig. 2B shows some bending in very short spindles, <∼3 μm, this is an artifact of the difficulty in accurately fitting curvature in short spindles.) We also quantify spindle elongation in both cerulenin-treated and control cells. Interestingly, while both initially elongate at similar rates, spindle elongation in cerulenin-treated cells slows down compared to control spindles when they reach a length of ∼4-5 μm, the same length at which they tend to buckle (Figure 2C). These data suggest that nuclear envelope tension is at least capable of slowing spindle elongation, and that the force that it exerts when it does so is sufficient to bend the spindle.

**Figure 2.**
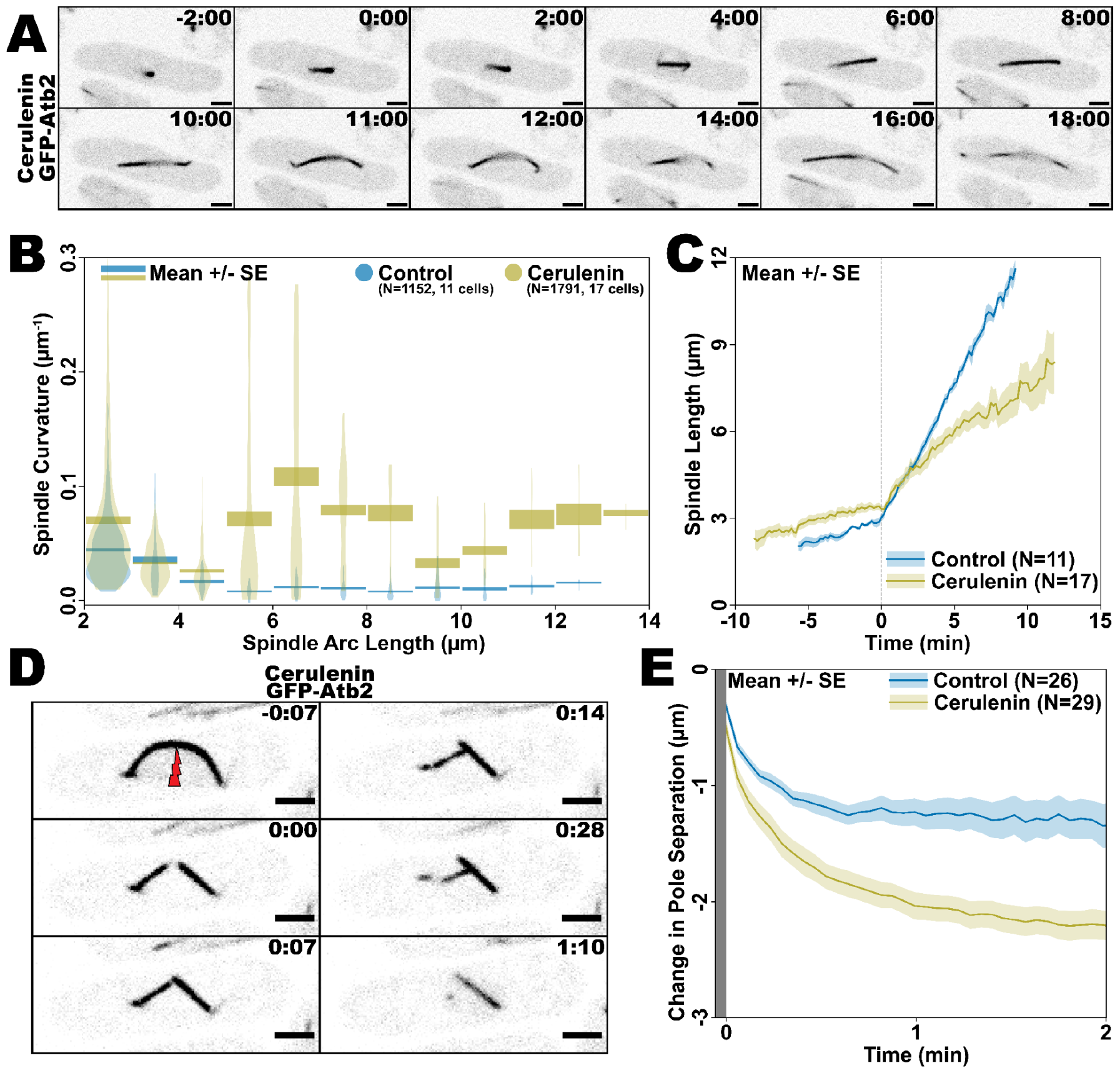
Increased nuclear envelope tension both bends spindles and slows their elongation. All scale bars, 2 μm, timestamps, min:sec.All cells strain MWE2. (A) Typical example of a cerulenin-treated *S. pombe* spindle bending as it elongates during anaphase (beginning around 10-11:00). (B) When elongating during anaphase, cerulenin-treated *S. pombe* spindles (gold) often begin bending when they are ∼5 μm in length, whereas untreated spindles rarely bend. The area of each violin corresponds to the number of data points in that bin, relative to the total number of points. Shaded region: mean +/-s.e.m. Outlying data (less than 0.44% of all data), with curvature values greater than 0.3 μm-1 are not shown here but are shown in Supp. Fig. 1. (C) Spindles in cerulenin-treated *S. pombe* (gold) elongate slower during anaphase than those in untreated cells (blue). Traces represent average change in pole separation, compared to the first frame after ablation, and shaded regions represent average +/-s.e.m. Traces aligned so that t=0 is the time when the rate of spindle elongation sharply increases. (D) Typical example of spindle ablation in cerulenin-treated *S. pombe* cell expressing GFP-Atb2. Dynamic changes to spindle structure seen here include spindle half straightening (0:00), spindle collapse, and unrepaired spindle half depolymerization (left half, 1:10). (E) Cerulenin-treated *S. pombe* spindles (gold) collapse much faster and more severely after ablation than ablated spindles in untreated *S. pombe* (blue). Traces represent average change in pole separation, compared to the last frame immediately prior to ablation, and shaded regions represent average +/-s.e.m.

To further verify that compressive force causes spindle buckling we selectively ablated mitotic spindles that appear curved (Figure 2D and Supplemental Video SV4). Consistent with inward forces on SPBs bending the spindle, the two spindle fragments straighten after they are ablated, when each half spindle now has room to fit within the nucleus. As with spindles in control *S. pombe*, the two spindle poles collapse toward each other after ablation in cerulenin-treated cells, however at a much greater rate (Figure 2E). In control *S. pombe* spindles, we have previously shown that pole collapse is a molecular motor-dependent response^13^. However, after severance of curved spindles, the incident angle between the two halves is much greater than that of ablated straight spindles (Figure 2D), a geometry that is likely to be less conducive to motor transport. Thus, we interpret the faster collapse in cerulenin-treated cells as likely due to an increase in compressive force acting on the spindle ends from the higher-tension nuclear envelope. Interestingly, and consistent with this interpretation, even though the two poles collapse toward each other, these spindles rarely reform a single microtubule bundle and, after failing to reconnect, ablated spindle halves often eventually depolymerize (Figure 2D, left spindle half).

### Release of nuclear envelope tension couples to spindle relaxation

While increased nuclear envelope tension could directly bend the spindle, other factors, such as chromosome entanglements or direct associations of the spindle with the nuclear envelope along its length, could also contribute. To further test these alternative possibilities, we performed experiments in which we ablated the nuclear envelope of cerulenin-treated *S. pombe* and tracked the changes to the spindle and nucleoplasm (Figure 3A and Supplemental Video SV5). Only in the case that nuclear envelope surface tension is compressing the spindle via the embedding of the SPBs would we expect ablation of the envelope far from the spindle itself to allow mechanical relaxation of the spindle. For these experiments, we use cells expressing both GFP-Atb2 (to visualize tubulin) and GFP-NLS (to verify nuclear leakage). In these experiments, we observe a viscoelastic-like response of the spindle. Usually within a minute following ablation, the spindle straightens from a curved conformation to its relaxed, straight state (Figure 3A, B). This implies a reduction in the compressive force acting on the spindle poles and provides additional evidence of nuclear envelope tension as the source of compression. Furthermore, we typically observe spindle relaxation in concurrence with nucleoplasm leakage, visualized by the weakening intensity of GFP-NLS signal in the nucleus (Figure 3A, B). This correlation is apparent at both the single cell and population levels (Figure 3A-C).

**Figure 3.**
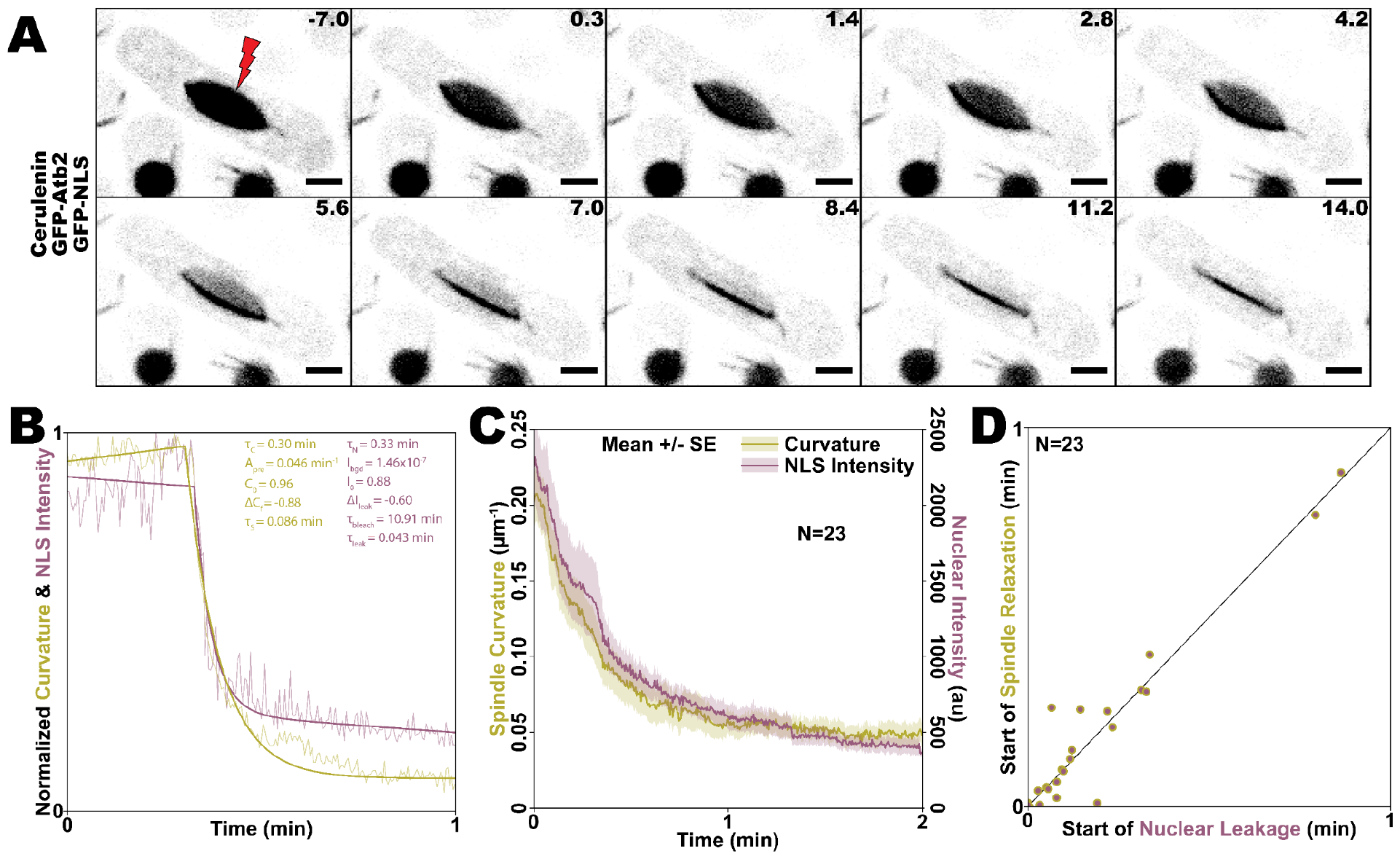
The fission yeast spindle and nuclear envelope are a mechanical pair. All cells strain MWE48. (A) Typical example of nuclear envelope ablation in cerulenin-treated *S. pombe* cell expressing GFP-Atb2 and GFP-NLS, showing nucleoplasm leakage and spindle relaxation, following nuclear envelope rupture. Scale bars, 2 μm. Timestamps, sec. (B) Example of typical NLS intensity (purple) and spindle curvature (gold) traces, following laser ablation of cerulenin-treated *S. pombe* nuclear envelopes. Fit (darker lines) and parameters (see Eq. 1 and 2) are given for each trace. (C) The averages of NLS intensity (purple) and spindle curvature (gold) show similar trajectories, following ablation. Traces represent averages across all videos and shaded regions represent average +/-s.e.m. (D) The initiations of nucleoplasm leakage and spindle relaxation, defined using best fit traces for each individual video (as in Figure 3B), are approximately simultaneous. Line shows 1:1 correlation.

Interestingly, at the individual cell level, we often observe a delay after ablation of the nuclear envelope before substantial spindle and nuclear envelope relaxation (Figure 3B). To quantify this behavior, we fit the curvature to the function

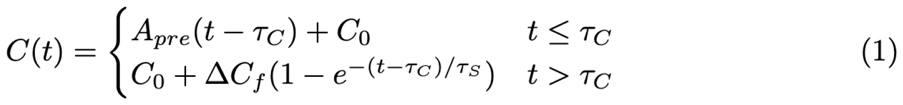

where *C*(*t*) is spindle curvature, *t* is time, and fit parameters are: *C*_0_, the curvature at the onset of straightening; Δ*C*_*f*_, the final change in curvature; τ_*C*_, the delay between ablation and straightening; τ_*s*_, the timescale of straightening; and *A*_*pre*_, a constant that takes into account variation in curvature before straightening begins. We quantified and fit the NLS intensity to:

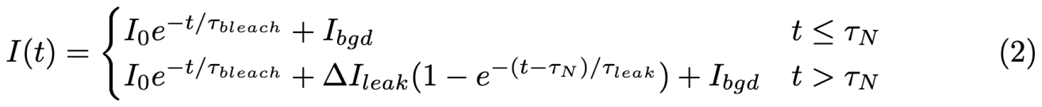

where *I*(*t*) is NLS intensity and fit parameters are: *I*_0_, the intensity at the onset of leakage; Δ*I*_*leak*_, the total change in intensity due to leakage; ΔI_*bleach*_, the additional change in intensity due to bleaching; τ_*N*_, the delay between ablation and leakage; τ_*leak*_, the timescale of leakage; τ_*bleac*h_the timescale of photobleaching; and I _*bgd*_, a constant that takes into account fluorescent background. When quantifying both curvature and intensity, we observe a delay between ablation and the cell’s response. These two delays in spindle straightening (τ_*C*_) and in nucleoplasm leakage onset (τ_*N*_) are highly correlated (Figure 3D), suggesting a causative relationship. In other words, we infer that the onset of leakage indicates mechanical failure of the nuclear envelope, resulting in a drop of envelope tension, which in turn allows the spindle to relax.

### The fission yeast spindle and nuclear envelope are a mechanical pair

Lastly, we build a simple mechanical model that quantitatively relates the behaviors of the spindle and nuclear envelope. First, we use the experimental results from this paper along with previous work to describe the relationship between nuclear envelope surface tension and spindle buckling. In general, buckling of a solid occurs above a particular threshold force. In this case, we assume that the nuclear envelope applies this buckling force on the spindle via the spindle pole bodies, which connect to the spindle and are embedded in the envelope (Figure 4A). Using the spindle buckling model from Ward et. al.^34^, the force needed to buckle spindles is

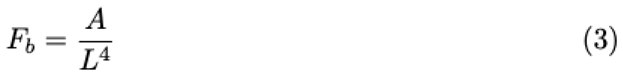

where the prefactor *A* = 1. 94 × 10^4^ *pN* µ*m*^4^ for wild-type fission yeast spindles. Consistent with Fig. 2A, which shows that spindles only bend above a threshold length, this model predicts a decreasing buckling force as spindle length extends, with predicted *F*_*b*_ = 31 *pN* for 5 μm-long spindles^34^. Because distortion of the nuclear envelope requires that the envelope bend and stretch, this deformation is energetically unfavorable and requires force. Previous modeling work found the characteristic force scale associated with deforming a membrane as

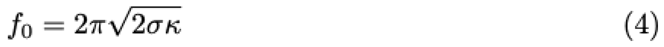

where σ and κ are the surface tension and bending rigidity of the nuclear envelope respectively^35–37^. Using measured values for wild-type *S. pombe*^*38,39*^ (see Supplementary Table) we estimate the force scale for distorting the fission yeast nuclear envelope as *f*_0_ = 6. 4 *pN*.

**Figure 4.**
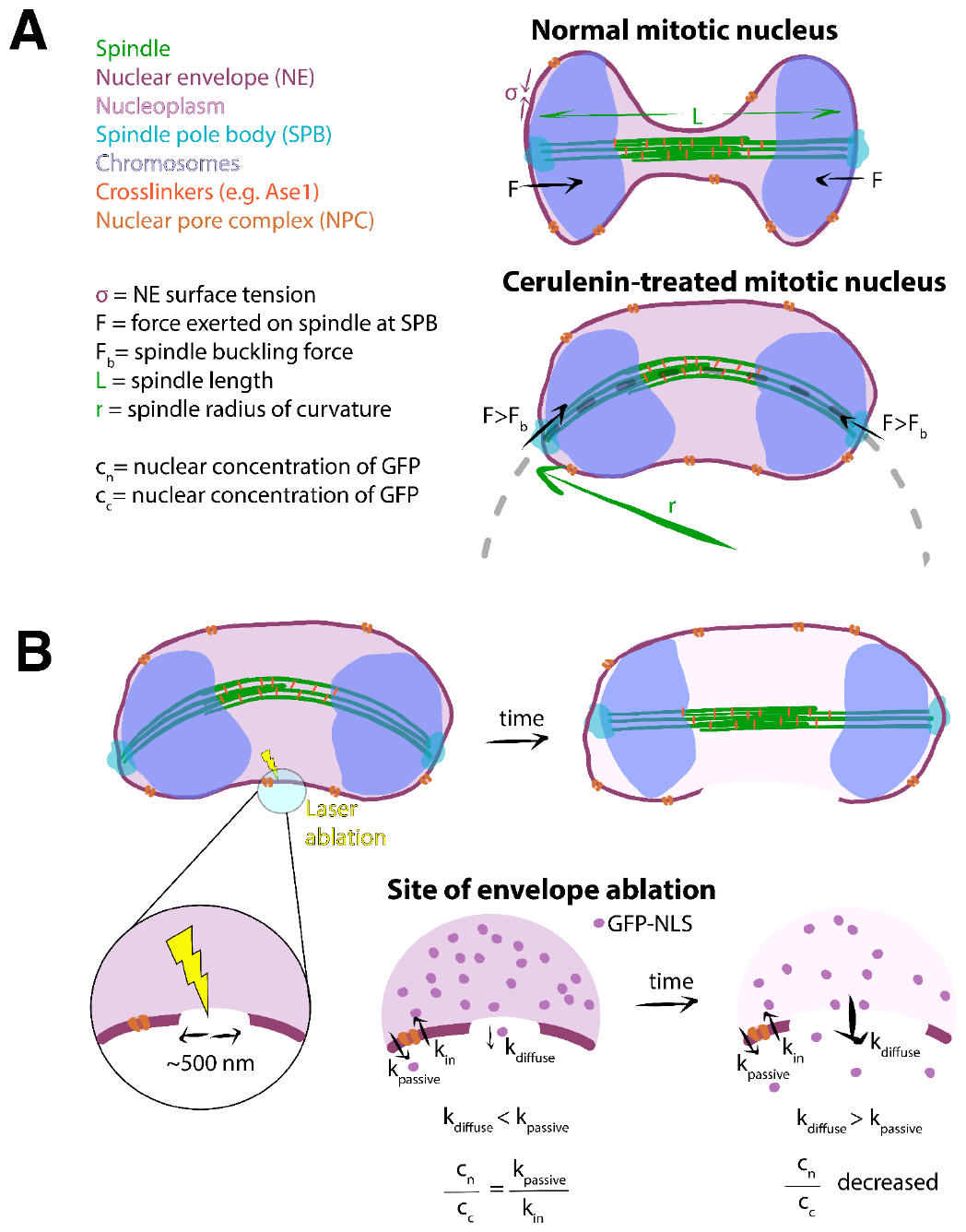
Mechanical model explaining how the nuclear envelope can shape the mitotic spindle. (A) During normal chromosome elongation, the *S. pombe* mitotic spindle elongates and is able to oppose the inward force on the SPBs, which results from the surface tension in the nuclear envelope. When cells are treated with cerulenin, the surface tension increases, and the inward force on the SPBs is great enough to cause the spindle to buckle. (B) Following laser ablation, a hole that is diffraction-limited in size is created in the nuclear envelope, allowing GFP-NLS to diffuse into the cytoplasm. Meanwhile, nuclear pore complexes continue to pump GFP-NLS from the cytoplasm back into the nucleus. These two processes are in a fast quasi-equilibrium. At later times, the hole ruptures, both allowing more diffusion of GFP-NLS and lowering the envelope tension, allowing the spindle to straighten.

Therefore, in unperturbed fission yeast, the force generated by the tendency of the envelope to resist deformation is unlikely to reach the *F*_*b*_ = 31 *pN* required to induce buckling of 5 μm-long spindles, and indeed, spindle bending is quite rare in unperturbed cells. Assuming that the bending rigidity of the spindle and the nuclear envelope do not change when cells are treated with cerulenin, the increase in surface tension induced by cerulenin treatment can be estimated using the buckling force. To generate enough force for buckling, we estimate that cerulenin causes a ∼20-fold increase in surface tension, with

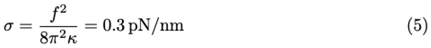

Second, we consider a quantitative description of the leakage of GFP-NLS out of the nucleus following laser ablation (Fig. 4B). We initially assumed that the drop in intensity we observe was due directly to GFP-NLS molecules diffusing out of the hole in the envelope generated by ablation. Previous work showed that escape of a diffusing particle through a round hole leads to an exponential decrease in concentration, with rate *k* = 1/τ = 4*Dr*/*V*, where *D* is the particle diffusion coefficient, *r* is the radius of the hole, and *V* is the volume of the nucleus^40^. To estimate the hole radius *r* = *V*/(4*D*τ), we measured *D* = 11 ± 5 µ*m*^2^ /*s* by FCS (Supp. Fig. 2), take τ=10 s as a typical value for the relaxation time constant (Fig. 3B-C), and estimate *V* ≈ 14 µ*m*^3^ (see Supplementary Table). Together, these predict a hole radius due to laser ablation of

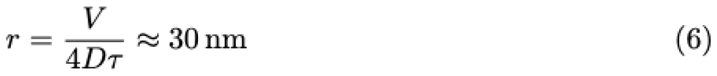

This estimate is unphysically small, given that damage due to laser ablation occurs in a diffraction-limited spot with a characteristic size ≈ 500 *nm*^41–43^. We thus consider additional mechanisms that may control the time course of both GFP-NLS concentration and spindle curvature.

Because GFP-NLS is actively imported into the nucleus and active import can occur rapidly, our data may reflect a balance between passive diffusion of GFP-NLS through the hole and active import as the hole size changes over time. We therefore construct a model that considers both of these effects (see Supplementary Discussion), yielding the following result:

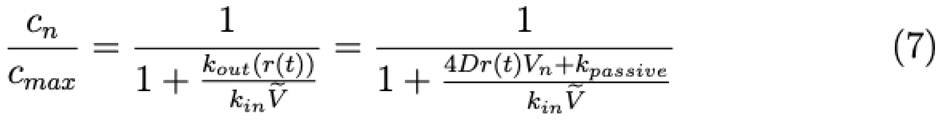

where *c*_*n*_ is the nuclear concentration of GFP-NLS, *c*_*max*_ is the concentration if all GFP-NLS molecules were in the nucleus, and 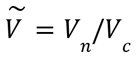 is the ratio of nuclear to cytoplasmic volume. The first-order rate constant of leakage *k*_*out*_ (*r*) = 4*Dr*(*t*)*V*_*n*_ + *k*_*passive*_ describes both diffusion through the hole in the nuclear envelope (which depends on the size of the hole) and ongoing passive leakage (through the nuclear-pore complex, assumed constant), while the first-order rate constant *k*_*in*_ describes active import through the nuclear-pore complex and is assumed constant. Before ablation, there is no hole and we predict 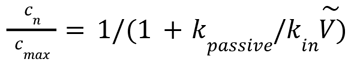 or, in terms of the more easily measurable cytoplasmic concentration,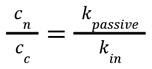. Using fluorescence intensity to measure the concentration ratio and taking reasonable estimates for diffusion rates (see Supplementary Table and Supplementary Discussion), we estimate that at a hole size of *r* ≈ 0.5 - 1.5 µ*m*, the nuclear intensity would be predicted to change due to significant diffusion of GFP-NLS out of the hole in the nuclear envelope. This micron-size hole is physically reasonable to observe after ablation.

Our model therefore explains why the drop in GFP-NLS intensity in the nucleus and the drop in spindle curvature have the same timescale. The decrease in intensity is likely limited not by the rate of diffusion through the hole, but by the hole size, which must therefore change over time. While the hole caused by ablation is likely initially small in area, it may trigger a subsequent catastrophic event in which the hole expands and nuclear envelope tension plummets, allowing the spindle to straighten. Such a response would explain the delay between ablation and both the beginning of nuclear leakage and the beginning of spindle straightening (Figure 3E): this delay indicates the time between ablation and envelope rupture to a sufficiently large hole size to compete with active GFP-NLS import. Additionally, the estimated size at which the hole allows a significant drop in GFP-NLS concentration, *r* ≈ 1 µ*m*, is physically consistent with the size of deformation we would expect to be required to allow the spindle to straighten. Determining what underlies the timescale of the rupture itself will be an interesting area of further investigation: one can imagine that it might depend on nuclear envelope material properties, larger scale rearrangements of the cytoplasm, nucleus, and spindle, or a combination of both.

In total, our results demonstrate significant mechanical coupling, during mitosis, between the mitotic spindle and nuclear envelope in *S. pombe*. The rapid spindle collapse following spindle ablation in cerulenin-treated cells suggests the presence of a compressive force on the spindle poles, resulting from increased nuclear envelope tension. Likewise, the simultaneous nuclear envelope rupture and spindle relaxation after nuclear envelope ablation implies this compressive force can be lessened by relieving tension in the nuclear envelope. The high degree of crosslinking in the *S. pombe* spindle likely allows it to oppose a typical level of nuclear envelope tension to remain straight during normal mitosis, when nuclear envelope expansion accommodates an elongating anaphase spindle. However, even this level of mechanical reinforcement is insufficient to prevent bending when compression is increased by treatment with cerulenin. Future work could provide further insight into how the mechanical properties of these two functionally integrated structures, the mitotic spindle and the nuclear envelope, may have co-evolved to support robust chromosome segregation and genetic integrity.

## Materials and Methods

### RESOURCE AVAILABILITY

#### Lead contact

Further information and requests for resources and reagents should be directed to and will be fulfilled by the Lead Contact, Mary Williard Elting.

#### Materials availability

Fission yeast strains developed in this study are stored at NC State University and samples are available on request.

#### Data and code availability

The raw data reported in this study consist of microscopy data, which will be shared by the lead contact upon request, along with any additional information required to reanalyze the data.

### FISSION YEAST STRAINS AND CULTURE

For strain details, see Key Resources table. *S. pombe* strains MWE2 and MWE3 are from Fred Chang lab stock (original strains FC2861 and FC1984^25^, respectively). MWE48 was created by crossing *S. pombe* strains MWE2 and MWE40, which was from Gautam Dey lab stock (original strain GD208^44^). Cross was performed by tetrad dissection using standard methods^45^. All strains were cultured at 25C on YE5S plates using standard techniques^45^. For imaging, liquid cultures were grown in YE5S media at 25 °C with shaking by a rotating drum for 12-24 hours before imaging. To ensure that cells were in growth phase for imaging, we measured OD595 with a target of 0.1-0.2. If cells had grown beyond this point, we diluted them and allowed them to recover for ∼1 hour before imaging. As a method of increasing nuclear envelope tension in *S. pombe*, cells were treated with 0.3-1mM cerulenin for between 1 and 4 hours before imaging, from stock solutions at 50 mM in DMSO, as previously described^1,32^.

### LIVE CELL IMAGING, LASER ABLATION, AND FLUORESCENCE CORRELATION SPECTROSCOPY

Prior to imaging, samples were placed onto gelatin pads on microscope glass slides. For gelatin pads, 125 mg gelatin was added to 500 μL EMM5S and heated, in a tabletop dry heat bath at 90oC for at least 20 min. A small sample volume (∼5 μL) of the gelatin mixture was pipetted onto each slide, covered with a coverslip, and given a minimum of 30 min to solidify. For each microscope slide, 1 mL volume of cells suspended in YE5S liquid growth media were centrifuged (enough to see a pellet), using a tabletop centrifuge. Nearly all the supernatant was decanted and the cells were resuspended in the remaining supernatant. Next, 2 μL of resuspended cells were pipetted onto the center of the gelatin pad, which was immediately covered with a cover slip. Finally, the coverslip was sealed using VALAP (1:1:1: Vaseline:lanolin:paraffin). All samples, sealed between the gelatin pads and coverslips, were imaged at room temperature (∼22°C).

Spinning disk confocal live imaging and laser ablation experiments were performed similar to those described previously^10,13,15^. Live videos were captured using a Nikon Ti-E stand on an Andor Dragonfly spinning disk confocal fluorescence microscope; spinning disk dichroic Chroma ZT405/488/561/640rpc; 488 nm (50 mW) diode laser (240 ms exposures) with Borealis attachment (Andor); emission filter Chroma Chroma ET525/50m; and an Andor iXon3 camera. Imaging was performed with a 100x 1.45 Ph3 Nikon objective and a 1.5x magnifier (built-in to the Dragonfly system). An Andor Micropoint attachment with galvo-controlled steering was used for targeted laser ablation, delivering 20-30 3 ns pulses of 551 nm light at 20 Hz. Each cell was imaged until either spindle repair had been completed or the spindle had failed to repair after ∼5 min. For spindle ablation videos frames were collected every 3.5 s, while frames were collected every 280 ms for nuclear envelope ablation videos. Andor Fusion software was used to manage imaging and Andor IQ software was used to simultaneously manipulate the laser ablation system. We used GFP as a marker in all cases due to greater robustness against photobleaching when we ablate. Due to differences in localization, we were able to discriminate between the GFP-Atb2 and GFP-NLS signals.

Fluorescence correlation spectroscopy (FCS)^46–48^ measurements were performed with a time-resolved confocal fluorescence microscope (MicroTime 200, PicoQuant) with SymPhoTime64 software for data acquisition and analysis. A picosecond pulsed diode laser (485 nm) controlled by a laser driver module (SEPIA II) was used to excite the sample with a 60x 1.2 numerical aperture water immersion lens (Olympus UPlanSApo, Superachromat). A fast galvo beam scanning module (FLIMbee) was used for laser beam scanning. Time-correlated single photon counting (TCSPC) with a multichannel event timer (MultiHarp 150) in time-tagged time-resolved (TTTR) measurement mode applied time tags to individual photons detected at a single photon avalanche diode (SPAD) with a 511/20 bandpass filter. Detected photons were also marked with a location in the beam scan of the sample to enable fluorescence lifetime imaging microscopy (FLIM). A FLIM image was acquired and the nuclei of cells were selectively pre-bleached by scanning with an average laser power density of ∼ 3 *kW*/*cm*^2^. FCS measurements were acquired by point clicking on a bleached location within the nucleus and collecting photons from diffusing GFP-NLS molecules. An autocorrelation was performed on the fluorescence intensity trace and the decay fitted to a pure diffusion model (see Supplementary Figure S2) to extract a diffusion time. The diffusion coefficient was calculated by characterizing the effective confocal volume (∼ 1 femtoliter) and kappa value (∼ 6) with several fluorescent dyes with a known diffusion coefficient. Simultaneous detection of the fluorescence lifetime of the diffusing species (∼2.7 ns) was consistent with GFP.

### QUANTIFICATION AND STATISTICAL ANALYSIS

#### Image and video preparation and editing

To optimize the identification and tracking of spindle and nuclear envelope features, modifications were made to fluorescence microscopy images and videos using FIJI^49^. First, images and videos were cropped to show only cells of interest and extra frames were eliminated. Typically, linear adjustments were made to the brightness and contrast of the images, in order to track features more clearly. For measurements of NLS intensity, however, pixel intensities were measured from unadjusted images. For immunofluorescence images, the same brightness and contrast scaling was used for all images in each set.

#### Tracking of spindle features in ablation videos

All quantitative data regarding post-ablation spindle dynamics was collected via a tracking program, home-written in Python. For each ablated spindle, the two spindle poles were tracked following ablation, using this program, and the ‘line’ tool in FIJI was used to measure the length of each spindle immediately prior to ablation (Supplemental Figure 3, A and B). These data were then used to calculate the change in pole separation (length) for each spindle over time, following ablation (Supplemental Figure 3C). Additionally, the positions of the two new plus-ends of each ablated spindle half were tracked throughout the video. Our tracking program includes a method for indicating whether or not spindle repair has occurred, with the reconnection of the two ablated spindle halves, in each frame. The data for frames collected before the reformation of a single spindle was used to compute time traces for the change in spindle half length (Supplemental Figure 3D).

We define spindle half splaying as the lateral dissociation of microtubule ends from each other in ablated spindle halves. Splay state is determined by eye in FIJI, using videos in which the brightness and contrast have been linearly adjusted for clarity. If a spindle half appears splayed in any frame of a video, that spindle half is counted as splaying. Otherwise, it is counted as a spindle that does not splay. For videos in which splay state is not readily apparent throughout, for one or both spindle halves, no splaying designation is made for the unclear half/halves. In some cases, a spindle half depolymerizes within the first few minutes following ablation and is therefore not included in our splay state analysis.

#### Quantification of spindle relaxation and nucleoplasm leakage

For all nuclear envelope ablation videos, data was collected on the time-evolution of spindle curvature using a home-written Matlab program. The program takes microscope image files and fits a quadratic function to a chosen object in the image. It then outputs curvature and length data from that fitted curve. For videos this process was semi-automated to perform the fit frame-by-frame.

Another program, home-written in Python was used to track the rate of nucleoplasm leakage from the nucleus of each cell following nuclear envelope ablation. This program requires the unadjusted video, spindle curvature data, and spindle length data as inputs. Using these inputs, nuclear intensity is calculated for each frame as the average GFP intensity (after background subtraction) of a 25-pixel square near the center of the nucleus. Both NLS and spindle curvature traces are normalized to their highest value for that video. This same program is then used to compute least squares fit curves for both post-ablation spindle curvature and nuclear intensity using the equations given in the main text.

### KEY RESOURCES TABLE

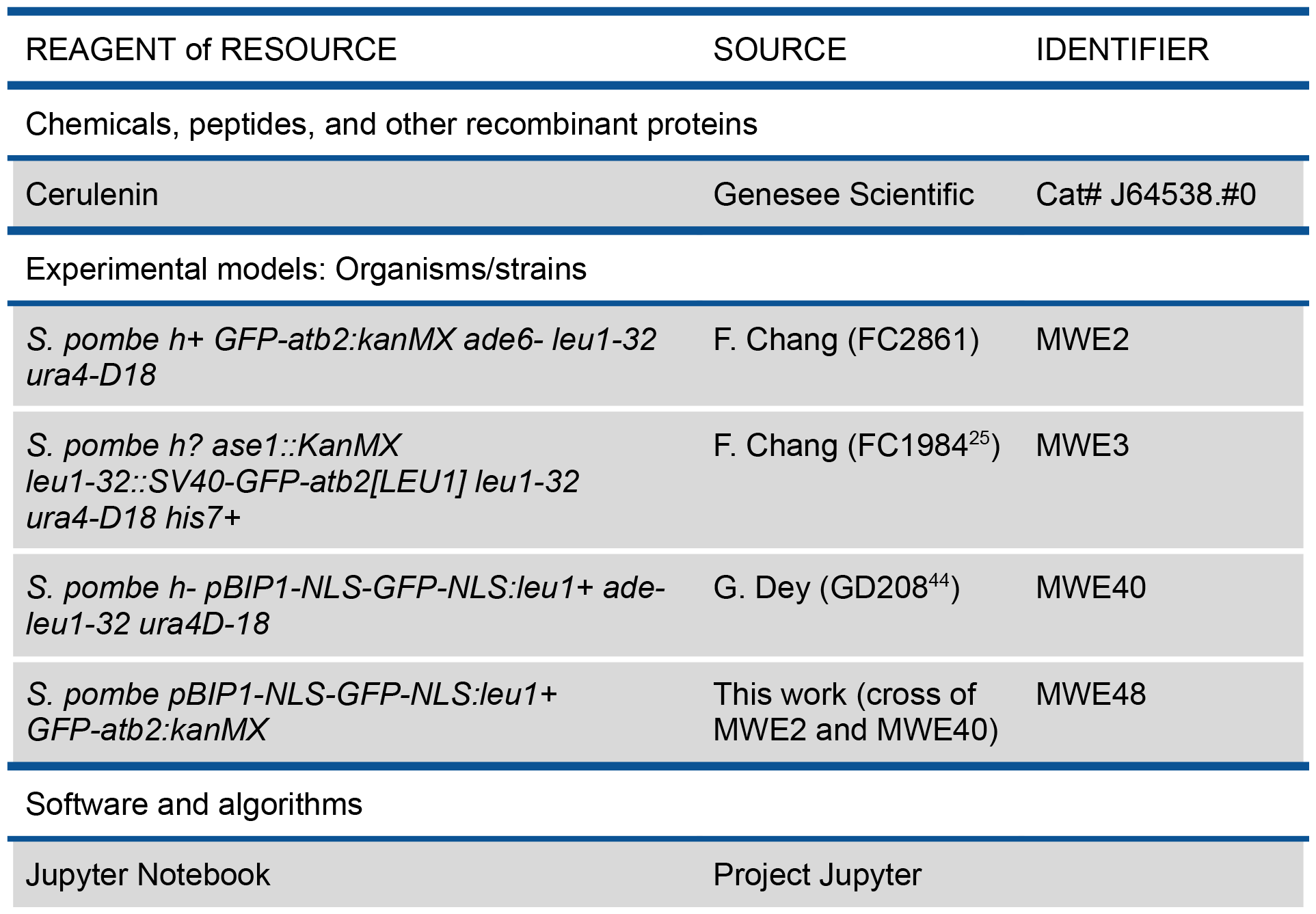
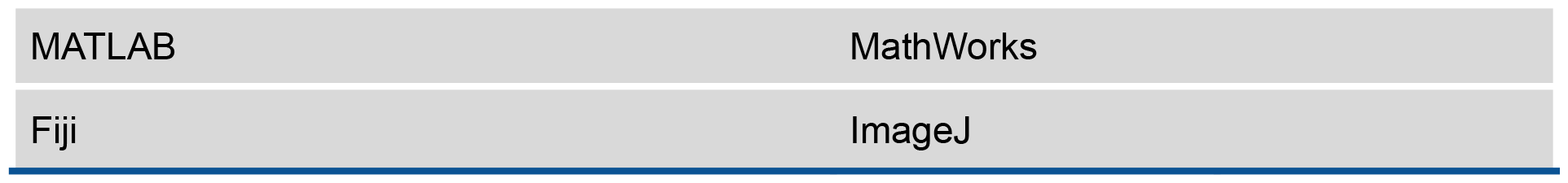

| REAGENT of RESOURCE | SOURCE | IDENTIFIER |
| --- | --- | --- |
| Chemicals, peptides, and other recombinant proteins |  |  |
| Cerulenin | Genesee Scientific | Cat# J64538.#0 |
| Experimental models: Organisms/strains |  |  |
| *S. pombe h+ GFP-atb2:kanMX ade6- leu1-32 ura4-D18* | F. Chang (FC2861) | MWE2 |
| *S. pombe h? ase1::KanMX leu1-32::SV40-GFP-atb2[LEU1] leu1-32 ura4-D18 his7+* | F. Chang (FC1984<sup>25</sup>) | MWE3 |
| *S. pombe h- pBIP1-NLS-GFP-NLS:leu1+ ade-leu1-32 ura4D-18* | G. Dey (GD208<sup>44</sup>) | MWE40 |
| *S. pombe pBIP1-NLS-GFP-NLS:leu1+ GFP-atb2:kanMX* | This work (cross of MWE2 and MWE40) | MWE48 |
| Software and algorithms |  |  |
| Jupyter Notebook | Project Jupyter |  |

|  |  |
| --- | --- |
| MATLAB | MathWorks |
| Fiji | ImageJ |

## Supporting information

Vidoe SV4

## ACKNOWLEDGEMENTS

We thank G. Dey, F. Chang, A. Molînes, S. Oliferenko, and members of the Elting lab, particularly Reem Hakeem, for advice and helpful discussions, and we thank F. Chang, G. Dey, and S. Oliferenko for strains. We thank the Weninger Lab (NCSU) and Wang Lab (NCSU) for sharing lab space and equipment. We thank the Cellular and Molecular Imaging Facility (CMIF) at NCSU for microscopy support. This work was supported by NIH 1R35GM138083 and NSF 2133276.

