## Supplemental Text, Figures, and Table for "Mechanical coupling with the nuclear envelope shapes the *S. pombe* mitotic spindle"

### Supplementary Material

**Supplementary Video Legends**

**Supplemental Video SV1:** Typical example of an ablated spindle in an *S. pombe* cell expressing GFP-Atb2. Following ablation, the spindle collapses as the distance between the two spindle poles decreases. Eventually, the mechanical connection between the two ablated spindle halves is re-established, restoring the spindle to a single bundle.

**Supplemental Video SV2:** Typical example of an ablated spindle in an *S. pombe ase1Δ* cell expressing GFP-Atb2. Here, the two spindle halves depolymerize shortly after ablation.

**Supplemental Video SV3**: Typical example of an elongating spindle in an *S. pombe* cell expressing GFP-Atb2 and treated with cerulenin. The spindle is prone to bending, as it elongates, under the compressive force exerted on the spindle poles by the nuclear envelope (10:00-15:00).

**Supplemental Video SV4:** Typical example of an ablated curved spindle in an *S. pombe* cell expressing GFP-Atb2 and treated with cerulenin. Post-ablation spindle collapse is more rapid and severe than in untreated *S. pombe*. Commonly, ablated spindle halves will depolymerize, after failing to reform a spindle.

**Supplemental Video SV5:** Typical example of an ablated deformed nuclear envelope in an *S. pombe* cell expressing GFP-Atb2 and GFP-NLS and treated with cerulenin. Both nucleoplasm leakage and spindle relaxation can be seen, following ablation of the nuclear envelope distal to the curved spindle.

#### Supplementary Figures

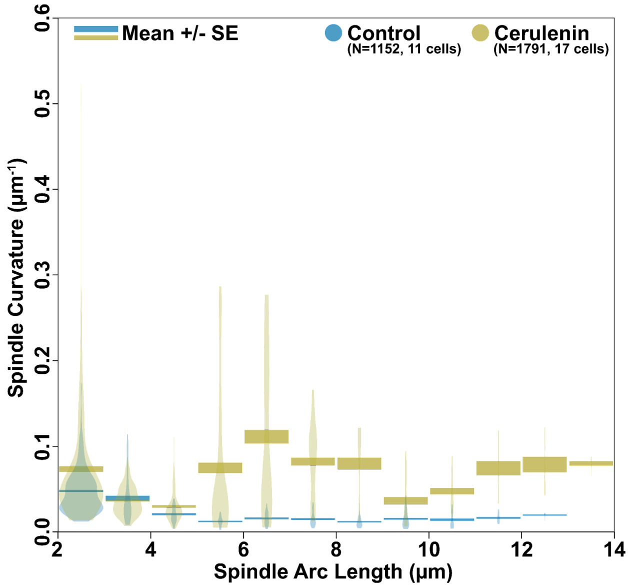


**Supplemental Figure 1**. Data from Figure 2B, with y-axis adjusted to show outlier data points (less than 0.44% of all data), which are not included in Figure 2B.

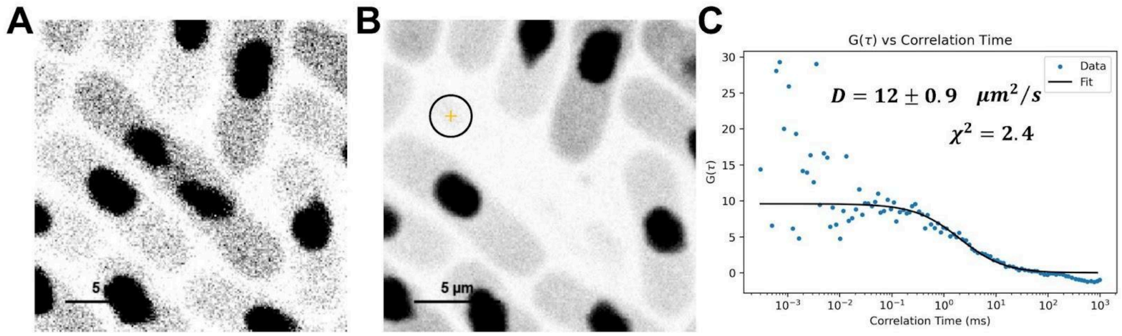


**Supplemental Figure 2**. Fluorescence correlation spectroscopy (FCS) demonstrates nuclear diffusion of GFP-NLS. (A) Pre-bleached image of *S. pombe* cell expressing GFP-NLS. Cells are strain MWE40. (B) Post-bleached image of *S. pombe* cell expressing GFP-NLS. The laser power and intensity scale for *B* is 10x compared to *A*. (C) FCS decay (scatter plot) from autocorrelation of fluorescence intensity trace collected at the location circled in *B*. The decay was fit to a pure diffusion model of the form $G(\tau) = \left(N \cdot (1 + \frac{\tau}{\tau_D}) \cdot \sqrt{1 + \frac{\tau}{\tau_D} \cdot \frac{\omega_0^2}{z_0^2}}\right)^{-1}$ (black line), where $\tau$ is a delay time, $N$ is the average number of molecules in the confocal volume, $\tau_D$ is the characteristic diffusion time, $\omega_0$ is the lateral beam radius, and $z_0$ is the axial beam radius. The diffusion coefficient, $D$, was calculated using the relation $D = \frac{\omega_0^2}{4\tau_D}$ after calibration of the effective confocal volume ($V_{eff}$) and kappa value ($k_{eff} = \frac{z_0}{\omega_0}$) with several dyes of known diffusion coefficient. Similar measurements were performed 12 times in 4 nuclei, resulting in a measured diffusion coefficient of $D = 11 \pm 5\ \mu m^2/s$ (mean +/- s.d.)

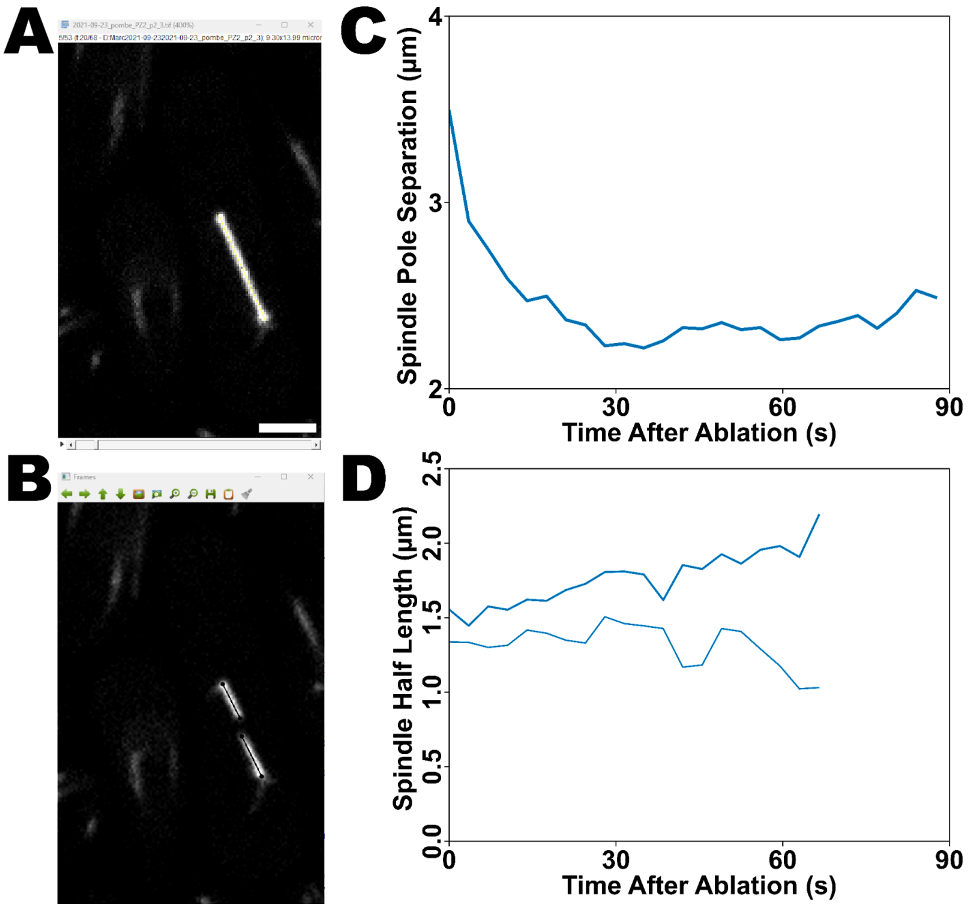


**Supplemental Figure 3.** (A) Screenshot showing the measurement of pre-ablation spindle length, using the line tool in Fiji. (B) Screenshot of a window from a home-written Python program used to label spindle poles and spindle half ablated plus-ends in the frames following ablation. (C) Example trace of post-ablation spindle pole separation for the spindle shown in panels A and B and Supplemental Video SV1. (D) Example trace of post-ablation spindle half length for the two ablated spindle halves shown in panel B and Supplemental Video SV1.

**Supplementary Table: Key parameters determined or used in the model**

| Variable name | Description | Relevant main text equation | Value | Source of value |
|---|---|---|---|---|
| $\tau_C$ | Delay between ablation and onset of straightening | 1 | 11 +/- 3 s | Fits to all cells as in Fig. 3B to Eq. 1 (mean +/- s.e.m.) |
| $\tau_S$ | Timescale of straightening (after straightening begins) | 1 | 12 +/- 2 s | Fits to all cells as in Fig. 3B to Eq. 1 (mean +/- s.e.m.) |
| $\tau_N$ | Delay between ablation and onset of nuclear leakage | 2 | 11 +/- 3 s | Fits to all cells as in Fig. 3B to Eq. 1 (mean +/- s.e.m.) |
| $\tau_{leak}$ | Time scale of leakage (after leakage begins) | 2 | 7 +/- 3 s | Fits to all cells as in Fig. 3B to Eq. 2 (mean +/- s.e.m.) |
| $A$ | Buckling prefactor | 3 | $1.94 \times 10^4 pN\, \mu m^4$ | Ref. [1] |
| $F_b$ | Buckling force | 3 | $31\, pN$ | Ref.[1] |
| $f_0$ | Force scale for nuclear envelope bending | 4 | $6.4\, pN$ | Ref.[2–4] and values below |
| σ | Normal nuclear envelope surface tension | 4 | $0.013\, pN/nm$ | Ref. [5,6] |
| κ | Nuclear envelope bending rigidity | 4, 5 | $40\, pN\, nm$ | Ref. [5,6] |
| $D$ | Diffusion coefficient of GFP-NLS in the nucleus | 6, 7 | $11 \pm 5\, \mu m^2/s$ | Fits to all cells as in Supp. Fig. 2C (mean +/- s.e.m.) |
| $V$ | Volume of the nucleus | 6, 7 | $14\, \mu m^3$ | $\frac{4}{3}\pi R^3$, $R \sim 1.5\, \mu m$ |
| $I_{nc}$ | Ratio of nuclear to cytoplasmic intensity of GFP-NLS | n/a | $23 \pm 6$ | Confocal fluorescence imaging of intensity in n=5 cells (mean +/- s.e.m.) |
| $\tau_{in}$ | Timescale of active import into the nucleus | n/a | 10-30 ms | Ref. [7–10] |
| $\tau_{passive}$ | Timescale of passive diffusion/leakage | n/a | 200-600 ms | $\tau_{passive} = \tau_{in} I_{nc}$ |

**Supplementary Discussion: Model of nuclear envelope leakage after ablation**

Because GFP-NLS is actively imported into the nucleus and active import can occur rapidly, our data may reflect a balance between passive diffusion of GFP-NLS through the hole and active import as the hole size changes over time. To model this, we suppose that there are $N_n(t)$ GFP-NLS molecules in the nucleus and $N_c(t)$ in the cytoplasm. We additionally assume that over the 1-minute timescale of the ablation experiments that the total number of GFP-NLS molecules does not significantly change, with a total number $N_0 = N_n + N_c$ and concentrations $c_n = N_n/V_n$, $c_c = N_c/V_c$, and $c_{max} = N_0/V_n$ (the maximum possible concentration if all molecules were in the nucleus), where $V_n$ is the nuclear volume and $V_c$ the cytoplasmic volume. The rate of change of the nuclear GFP-NLS concentration is

$$\frac{dc_n}{dt} = -k_{out}(r)c_n + k_{in}c_c \quad (1)$$

where the first-order rate constant of leakate $k_{out}(r) = 4Dr(t)V_n + k_{diffuse}$ describes both diffusion through the hole in the nuclear envelope (which depends on the size of the hole) and ongoing passive leakage (through the nuclear-pore complex, assumed constant), while the first-order rate constant $k_{in}$ describes active import through the nuclear-pore complex and is assumed constant. Substituting $N_c = N_o - N_n$ and the appropriate volume factors into Supp. Eq. (1) gives us

$$\frac{dc_n}{dt} = -(k_{out}(r) + k_{in}\widetilde{V})c_n + k_{in}\widetilde{V}c_{max} \quad (2)$$

where $\widetilde{V} = V_n/V_c$ is the ratio of nuclear to cytoplasmic volume. We treat Supp. Eq. (2) quasistatically, assuming that the import and export rates are rapid compared to the tens of seconds required for full relaxation of the GFP-NLS intensity. Therefore, we set the time derivative term on the left side of Supp. Eq. (2) to zero, and solve for the fraction of total molecules that are in the nucleus

$$\frac{c_n}{c_{max}} = \frac{1}{1+\frac{k_{out}(r(t))}{k_{in}\widetilde{V}}} = \frac{1}{1+\frac{4Dr(t)V_n+k_{passive}}{k_{in}\widetilde{V}}} \quad (3)$$

Before ablation, there is no hole and we predict $\frac{c_n}{c_{max}} = 1/(1 + k_{passive}/k_{in}\widetilde{V})$, or, in terms of the more easily measurable cytoplasmic concentration, $\frac{c_n}{c_c} = \frac{k_{passive}}{k_{in}}$. Since average fluorescence intensity is proportional to concentration, we measured the average fluorescence intensity of GFP-NLS in the nucleus relative to the cytoplasm $I_{nc}$ and found an average value $I_{nc} = c_n/c_c = k_{in}/k_{passive} \approx 23 \pm 6$ (n=5 cells). This result shows that the active nuclear import rate is significantly higher than the passive export rate through the nuclear pore complex, as expected. Even once the hole is present, we expect that diffusion through the hole is initially slow compared to transport rates through the nuclear pore, meaning that $4DrV_n \ll k_{passive}$, so the denominator of Supp. Eq. (3) does not initially increase. This could explain the initial delay observed in traces as in Fig. 3B, where the GFP-NLS concentration is initially approximately constant. The GFP-NLS concentration will first begin to show a noticeable decrease when the rate of GFP-NLS leakage through the ablated hole is comparable to the rate of passive nuclear export, meaning $4Dr(t)V_n \approx k_{passive}$. To estimate the hole size where this crossover occurs, we estimated $k_{in}$ and $k_{passive}$. Measurements of active transport through the nuclear pore complex find very short residence times of transported particles in the pore, in the range of 1-10 ms[7,8] and that the timescale of nuclear transport is typically limited by the time required for molecules to diffuse to a nuclear pore complex[10]. In fission yeast, nuclear pore complexes are typically separated by about 0.4 $\mu m$[9], making this a typical distance that a GFP-NLS must diffuse to reach a nuclear pore complex and re-enter the nucleus. This diffusive timescale for GFP-NLS is approximately $\tau_D \approx l^2/D \approx 15$ ms. This suggests that the timescale of active import is in the range of $\tau_{in}$ ≈10-30 ms, including both diffusion and transport time (reflecting the variability in transport rate[11]). The timescale for passive export is approximately 20 times larger, based on our intensity ratio estimate. Therefore we estimate the export timescale $\tau_{passive} = 1/k_{passive} \approx$ 200-600 ms. Based on the prediction of our model that noticeable GFP-NLS diffusion through the hole will occur once $4Dr(t)V_n \approx k_{passive}$, we therefore estimate that the hole size must grow to a radius $r = V_n/(4D\tau_{passive})$ before the intensity decrease is observable. This gives an estimated range of hole size $r \approx$ 0.5 - 1.5 $\mu m$ at which the nuclear intensity would be predicted to change due to significant diffusion of GFP-NLS out of the hole in the nuclear envelope. This micron-size hole is physically reasonable to observe after ablation.
